# NetLCP: An R package for prioritizing combinations of regulatory elements in the heterogeneous network with variant ‘switches’ detection

**DOI:** 10.1101/2022.10.06.511229

**Authors:** Mingyu Ran, Jiankai Xu

## Abstract

Numerous independent networks of regulatory elements, including lncRNA, circRNA and pathway, have been developed to crucial roles in computational systems biology. Crosstalks among those networks as a bridge to build and decode heterogeneous networks from multidimensional biological knowledge, aids to highlight regulatory elements. And combinations of regulatory elements (CREs) in the local area of heterogeneous network have been a hot issue due to its crucial role in biological processes. We introduce NetLCP, an R package with command and shiny-based GUI modes, for prioritizing CREs with variant ‘switches’ detection.

**Availability and implementation:** The NetLCP package and documentations are freely available at https://github.com/mortyran/NetLCP.

**Supplementary information:** Supplementary data are available online

## 1 Introduction

With recent volume escalation in high-throughput data, networks of regulatory elements, including lncRNA, circRNA and pathway, have sprung up and are becoming an indispensable issue of computational systems biology (Li *et al*., 2020; Yang and Li, 2021). Constructing heterogeneous network from multidimensional biological knowledge is an effective way to deal with deficient information in single networks (Ding *et al*., 2020; Yang *et al*., 2022). Combinations of regulatory elements (CREs) in the local area of heterogeneous network, such as lncRNA-miRNA-mRNA, have been widely demonstrated to involve in various biological processes. As the genes with eQTL effects are more tolerant to loss-of-function variants in their coding region (Võsa *et al*., 2021), sufficient eQTLs in CREs could improve transcriptional stability to maintain several functions. Besides, variants on the binding sites of genes can lead to dysfunction of CREs as ‘switches’ (Moszyńska *et al*., 2017). Nevertheless, manual implementation inevitably confronts diverse obstacles, particularly in processing and integrating biological networks, considering their volume, redundancy and complexity.

In this work, we constructed five heterogeneous networks consisting of seven types of independent networks which were built on current popular methods (**Supplementary Materials S2**). The crosstalks among the independent networks were mainly derived from experimentally verified biological regulatory data. A state-of-art diffusion algorithm, RWR-MH, was embedded to highlight regulatory elements by their similarities with input miRNA/mRNA in the same low-dimensional vector space (Valdeolivas *et al*., 2019; Ran *et al*., 2021). Because of high tolerance of the genes with eQTL effects to loss-of-function variants, we integrated eQTLs into CREs and further prioritized CREs by the number of their eQTLs. NetLCP provides advices about regulatory elements or CREs which are associated with user-interested biological processes or functions.

## 2 Implementation

The whole workflow of NetLCP is illustrated in **Figure 1**. A large heterogeneous network comprised a background network (regulatory element similarity network) and two input networks (miRNA and mRNA similarity network). Five such heterogeneous networks were constructed. The regulatory elements contained lncRNA, circRNA, KEGGPath, ReactomePath and WikipathwayPath. Details of computational methods can be found in **Supplementary Materials S2**. Biological regulations were mainly derived from experimentally supported data. NetLCP highlighted regulatory elements by measuring the similarities with low-dimensional vectors of input miRNA/mRNA, which were derived by decreasing the noise of heterogeneous networks using dimension reduction-equipped diffusion algorithm. For further prioritizing CREs in local area of heterogeneous network, we then integrated eQTLs of genes into CREs and performed statistics. Variants on the binding sites of genes were also mapped to CREs to detect potential variant ‘switches’-mediated dysfunction of prioritized CREs. Corresponding network visualization and statistics were provided. Data preparation in detail was elaborated in **Supplementary Materials S1**. We presented an application on cellular senescence-associated prioritization in **Supplementary Materials S3**.

**Figure 1.**
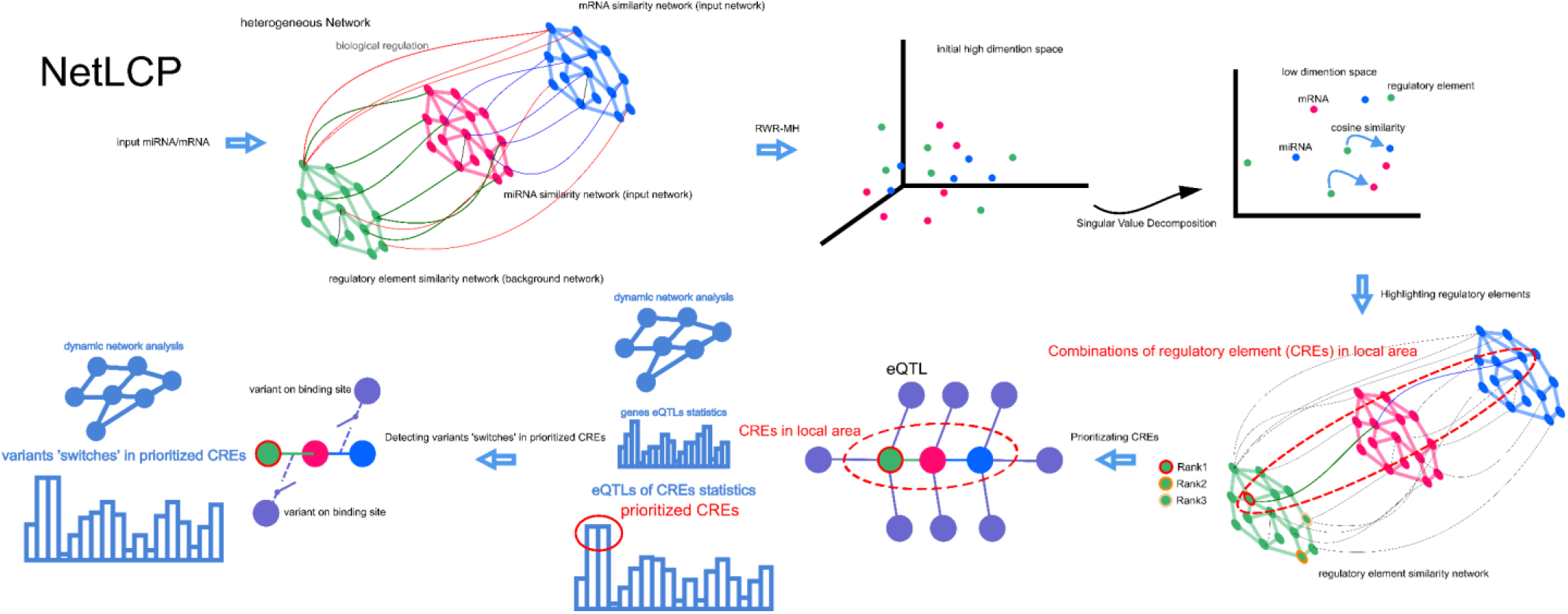
The workflow of NetLCP.

### 2.2 Highlighting regulatory elements in the heterogeneous network

NetLCP highlights regulatory elements (lncRNA, circRNA, KEGGPath, ReactomePath and WikipathwayPath) in the heterogeneous network, which have similar biological functions to the given input transcriptome (miRNA/mRNA). Regulatory elements are ranked by the similarity score. NetLCP produces a tab-delimited text files which records the prioritized elements with column names of lncRNA/circRNA/pathway ID, FunScore, OfficialName and Empirical P-value (alternative).

### 2.3 Prioritizing CREs

Current CREs in the local area of heterogeneous network contain circRNA-miRNA, lncRNA-miRNA, lncRNA-mRNA, miRNA-mRNA, miRNA-pathway, mRNA-pathway, lncRNA/circRNA-miRNA-mRNA, lncRNA/circRNA-miRNA-mRNA-pathway and miRNA-pathway. NetLCP prioritizes CREs by performing visual statistics on their eQTLs. Complete information of eQTLs including variant type, organ source (human blood/cancer type), location and data source can be exported.

### 2.4 Detecting variant ‘switches’ of regulatory elements

Variants on the binding sites of genes in CREs are composed of single nucleotide polymorphisms (SNPs) in lncRNA, 3’ untranslated region of mRNA and seed region of miRNA. The information, including element, variant type, variant location, and data source, can be output. Dynamic network visualization reveals the variant ‘switches’ in user interested CREs.

## 3 Conclusion

In conclusion, NetLCP prioritizes CREs by highlighting regulatory elements and detecting regulatory ‘switches’ in the heterogeneous network. By leveraging multidimensional biological knowledge, it provides a meaningful perspective on user-interested biological processes or functions.

