## Supplementary Materials for "NetLCP: An R package for prioritizing combinations of regulatory elements in the heterogeneous network with variant ‘switches’ detection"

**S1 Data Collection**

**1.1 Data statistics**

**1.2 Reference database**

**S2 Heterogeneous network construction**

**2.1 circRNA similarity network**

**2.1.1 Gaussian interaction profile kernel**

**2.1.2 circRNA sequence similarity**

**2.1.3 Merging**

**2.2 lncRNA similarity network**

**2.2.1 Literature expression network**

**2.2.2 lncRNA sequence similarity**

**2.2.3 Merging**

**2.3 Pathway similarity network**

**2.4 miRNA similarity network**

**2.5 mRNA similarity network**

**2.6 Highlighting regulatory elements in heterogeneous network**

**S3 Application**

**3.1 Highlighting regulatory elements**

**3.2 Prioritizing CREs**

**3.3 Detecting variant 'switches' in the top prioritized CREs**

**3.4 Application with NetLCP GUI**

#### S4 Tutorial

#### S1 Data Collection

##### 1.1 Data statistics

Experimentally supported regulation data of miRNA-mRNA, miRNA-lncRNA, miRNA-circRNA and lncRNA-mRNA were obtained, covering 1233555 miRNA-mRNA regulation pairs from ENCORI, NPInter, RAIN, miRNet, TarBase, miRTarBase, RISE and RNAInter, 84781 miRNA-lncRNA regulation pairs from LncBase, ENCORI, NPInter, RNAInter, TarBase, RAIN, miRNet and lncRNASNP2, 326845 miRNA-circRNA regulation pairs from ENCORI, NPInter, RNAInter and RAIN, 228323 lncRNA-mRNA regulation pairs from ENCORI, NPInter, RNAInter, RAIN and RISE.

Totally 5618406 cis-eQTLs, 15105 trans-eQTLs of mRNAs from eQTLGen and 4245785 cis-eQTLs, 5170 trans-eQTLs from ncRNA-eQTLs were obtained. 40844 CNVs, 505 COMIC somatic variants, 2399 SNPs and 617 TCGA somatic variants which might influence the ceRNA regulation were gained from LnCeVar. SNPs on the binding sites were categorized into seed SNP (SNPs on the seed region of miRNA) and UTR3 SNP (SNPs on the UTR3 of mRNA). 534067 seed SNPs and 2111524 UTR3 SNPs were collected from miRNASNPv3.

##### 1.2 Reference database

ENCORI: an open-source platform for studying the miRNA-ncRNA, miRNA-mRNA, ncRNA-RNA, RNA-RNA, RBP-ncRNA and RBP-mRNA interactions from CLIP-seq, degradome-seq and RNA-RNA interactome data. Here we extracted miRNA-mRNA, miRNA-lncRNA miRNA-circRNA and lncRNA-mRNA interaction data.

LncBase v3 (https://diana.e-ce.uth.gr/lncbasev3/home): a reference repository with experimentally supported miRNA targets on non-coding transcripts. Here we used miRNA-lncRNA regulatory data.

miRTarBase 2020: miRTarBase are manually collected by surveying pertinent literature after NLP of the text systematically to filter research articles related to functional studies of miRNAs. miRNA-mRNA, miRNA-lncRNA, miRNA-circRNA regulatory data were adopted.

TarBase v8: a reference database devoted to the indexing of experimentally supported microRNA (miRNA) targets. Here we adopted miRNA-mRNA, miRNA-lncRNA regulatory data.

RNAInter 2020: It updated of entries to >47 million by manually mining the literature and integrating six database resources with evidence from experimental and computational sources. Here we used only experimental sources in it.

LncRNASNP2: a database providing comprehensive resources of single nucleotide polymorphisms (SNPs) in human/mouse lncRNAs. It contained SNPs in lncRNAs, SNP effects on lncRNA structure, variant in lncRNAs and lncRNA-miRNA binding. Here we downloaded a part of this database including all experimentally validated miRNA-lncRNA interactions.

miRNet 2.0: miRNet is an easy-to-use, web-based platform designed to help elucidate microRNA (miRNA) functions by integrating users' data with existing knowledge via network-based visual analytics. Here we used experimentally supported miRNA-lncRNA, miRNA-mRNA regulatory data.

RAIN: a database of ncRNA–RNA and ncRNA–protein interactions and its integration with the STRING database of protein–protein interactions. Here we only adopted miRNA-mRNA, miRNA-lncRNA, lncRNA-mRNA and miRNA-circRNA regulatory data from experiments.

RISE: a comprehensive repository of RNA-RNA interactions. The interactions mainly come from recent transcriptome-wide sequencing-based experiments. Here we acquired miRNA-mRNA, miRNA-lncRNA and lncRNA-mRNA regulatory data.

NPInter v4.0: NPInter documents functional interactions between noncoding RNAs (except tRNAs and rRNAs) and biomolecules (proteins, RNAs and DNAs) which are experimentally verified. Here we accessed miRNA-mRNA, miRNA-lncRNA, miRNA-circRNA and lncRNA-mRNA regulatory data.

eQTLGen: The eQTLGen Consortium has been set up to identify the downstream consequences of trait-related genetic variants.

ncRNA-eQTL (miRNA-eQTL and lncRNA-eQTL): miRNA-eQTL aims to comprehensively provide miRNA related cis-eQTLs (SNPs affect local miRNA gene expression) and trans-eQTLs (SNPs affect distant miRNA gene expression).

lncRNA-eQTL: lncRNA-eQTL aims to comprehensively provide lncRNA related cis-eQTLs (SNPs affect local lncRNA gene expression) and trans-eQTLs (SNPs affect distant lncRNA gene expression).

miRNASNPv3: miRNASNP is a comprehensive resource of human miRNA-related SNPs, including their effects on target gain and loss.

LnCeVar: LnCeVar is a comprehensive database that aims to infer genomic variations that disturb lncRNA-associated ceRNA regulation.

#### S2 Heterogeneous network construction

##### 2.1 circRNA similarity network

###### 2.1.1 Gaussian interaction profile kernel

CircRNA-disease interaction data were derived from the three manually curated and experimentally verified databases, Circ2Disease, CircR2Disease(v1) and CircR2Disease(v2). Then the circBase ID cross-references were downloaded from circBase and circBank, the MeSH (Medical Subject Headings 2022) Term file in XML format was derived from National Library of Medicine using Python. After merging circRNA-disease associations from 3 databases and removing redundancy, we further unified the circRNA names to circBase IDs, the disease names to MeSH IDs. Totally 2132 circRNA-disease associations were obtained.

Gaussian interaction profile (GIP) kernel similarity model assumes that circRNAs typically exhibit similar disease-associated patterns and it has been widely applied in bioinformatics analysis such as measuring drug similarity (Yan *et al.*, 2019). In our study, GIP model is mathematically defined as formulas:

$$K_{GIP}(cir_i, cir_j) = exp\left(-y_{cir}\left\|C_{cir_i} - C_{cir_j}\right\|^2\right)$$

The GIP kernel similarity between circRNA $i$ and circRNA $j$ is termed as $K_{GIP}(cir_i, cir_j)$, $C_{cir_i}$ and $C_{cir_j}$ are the binary vectors, indicating whether circRNA $i$ and circRNA $j$ are associated with diseases. To deal with the influence of the sparsity of circRNA-disease association matrix, the parameter $y_{cir}$ is set to control the kernel bandwidth, which can be computed by following formula:

$$y_{cir} = n_{cir} \sum_{i=1}^{n_{cir}} \left\| C_{cir_i} \right\|^2$$

where $n_{cir}$ represents the number of circRNAs.

##### 2.1.2 circRNA sequence similarity

From circBank, the latest version of circRNA sequences were acquired. Chao game representation (CGR) (Jeffrey, 1990), which has been widely applied in a variety of bioinformatics tasks (Bonidia *et al.*, 2022; Anitas, 2022; Millán Arias *et al.*, 2022; Paul *et al.*, 2022). It takes position information and nonlinear relationship into consideration to obtain specific vector representation of sequences. The algorithm completely conserves the original information of the sequence in the 2-Dimensional coordinate system (CGR plane, a unit square, see **Supplementary Fig.1 A**) and embeds the position information into a mapping called frequency matrix of CGR graph (FCGR). Mathematically, it is presented as formulas.

$$P_n = 0.5 * (P_{n-1} + V_n) \quad n = 1,2,3...N$$

Where $P_n$ is the position of the Nth base of circRNA sequence, $P_0$ is starting point ($P_0 = (0.5, 0.5)$ ). $V_n$ represents the corner coordinate of the unit square $V_A(0,0), V_C(1,0), V_G(1,1), V_U(0,1)$. Furthermore, the CGR graph is transformed into $2^k * 2^k$ squares, i.e. FCGR (see **Supplementary Fig.1 B**). For $k = 1 (FCGR_1)$, CGR graph can be divided four sub-square and each sub-square represents a monomer nucleotide. Analogously, each sub-square represents a dimer nucleotide in the $FCGR_2$. In this study, we chose $FCGR_3$ to construct the circRNA sequence landscape as previous study shows (Shu *et al.*, 2021), i.e. the $i$th circRNA can be represented by $CLS_i$ and each $CLS_i$ can be separated into $2^3 * 2^3$ grids.

Then we extracted information rooting in x-axis ($X_k$), y-axis direction ($Y_k$) and digital features ($Z_k$) from each grid to construct the feature vector of circRNA sequence. Grid can be represented by formulas:

$$Grid_k = (X_k, Y_k, Z_k)$$

$X_k$ and $Y_k$ are calculated by accumulating the abscissa point $x$ and ordinate point $y$ in $Grid_k$ respectively. $Z_k$ is obtained by normalizing the grid points to quantify potential sequence features:

$$Z_k = \frac{N_k - \frac{\sum_{m=1}^{G} N_m}{G}}{\sqrt{\frac{1}{G}\sum_{m=1}^{G}(N_k - \frac{\sum_{m-1}^{G} N_m}{G})^2}}$$

Where $N_k$ is the number of points in $Grid_k$, G was the number of grids. Along these lines, $CLS_i$ can be converted to a vector, $(Grid_1, Grid_2 ... Grid_k)$, $Vec_i$. Ultimately, the similarity between circRNA $a$ and circRNA $b$, is measured by calculating Pearson correlation between $Vec_a$ and $Vec_b$.

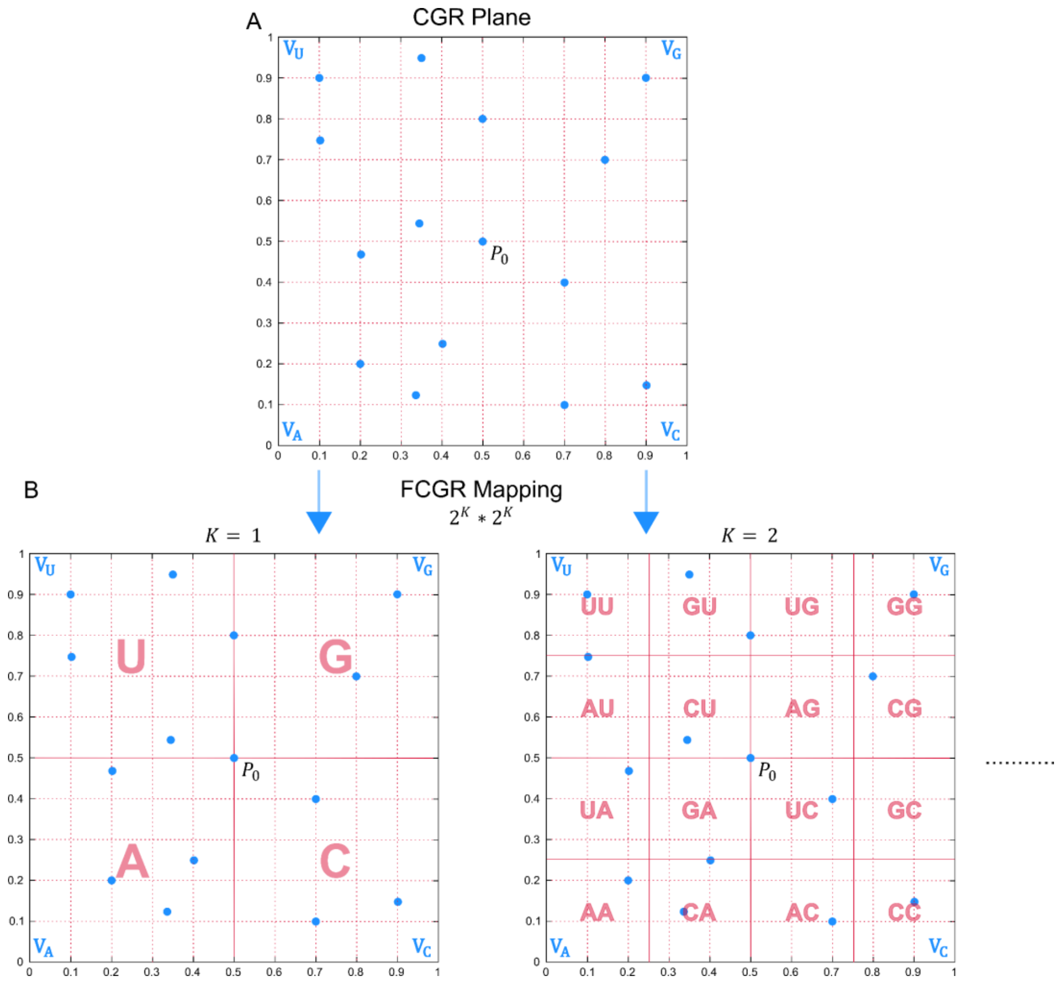


**Supplementary Fig.1. Workflow of generating circRNA sequence representation A) Sequence representation in CGR plane. B) Mapping of frequency matrix of CGR graph (FCGR)**

##### 2.1.3 Merging circRNA similarity network

We combined GIP- and CGR-based similarity by following formula:

$$\begin{cases} Sim = \alpha * GIP + \beta * CGR \\ \alpha + \beta = 1 \end{cases}$$

Here we assigned $\alpha$ to 0.5. Finally, 1156881 edges between 1519 circRNAs were obtained.

#### 2.2 lncRNA similarity network

##### 2.2.1 lncRNA functional similarity network

lncRNA functional similarity Network was constructed by the method of integrating four types of lncRNA similarity calculations including miRNA-, disease-, GTEx Expression-, and NONCODE Expression-based similarity measurements. The importance of each similarity measurement was characterized by the prediction power, which was represented by corresponding AUC value (Li *et al.*, 2020). In our research, for unifying the lncRNA names to Ensembl ID, we downloaded lncRNA name-mapping files from Ensembl and LNCipedia (v5.2) database.

##### 2.2.2 lncRNA sequence similarity

We downloaded lncRNA sequence data in latest version of Ensembl database. Sequence similarity was analogously measured by CGR method as described above (2.1.2).

##### 2.2.3 Merging lncRNA similarity network

We merged these two lncRNA similarity networks by the same method as described above (2.1.3). Finally, a large lncRNA similarity network containing 4530 lncRNAs and 7403280 edges was obtained.

#### 2.3 Pathway similarity network

Biological pathways comprise a cluster of interacting genes and can be regarded as a meaningful construct for inspecting the underlying biological mechanism of genes. Therefore, the similarity measurement of pathways can be naturally represented by the functional annotation similarity of pathway members. In this study, we adopted an information-theoretic definition of similarity in the natural language processing (NLP) area, which was based on comentropy theory. We first accessed the GENEONTOLOGY (http://geneontology.org/docs/go-annotations/) to gain the GO annotation file. Three reliable and comprehensive pathway resources, KEGG, Reactome and Wikipathway, were used for network construction. All the gene names were converted to Entrez ID. The similarity between given pathway $i$ and $j$ can be calculated by following formula:

$$sim(i,j) = \frac{2 * \sum_{t \in GO_i \cap GO_j} logP(t)}{\sum_{t \in GO_i} logP(t) + \sum_{t \in GO_j} logP(t)}$$

Where $t$ is a certain GO term, $GO_i$ and $GO_j$ represents the whole GO term set in pathway $i$ and $j$ respectively, $P(t)$ is the frequency of $t$ in the background GO term which was consist of the union of $GO_i$ and $GO_j$. Finally, three networks were constructed. There are 93642 edges between 334 nodes in KEGG pathway network, 2102150 edges between 2040 nodes in Reactome pathway network and 293250 edges between 703 nodes in Wikipathway pathway network respectively.

**2.4 miRNA similarity network**

MISIM v2 (http://www.lirmed.com/misim/) (Li *et al.*, 2019) infers miRNA functional similarity through integrating miRNA expression level and miRNA-disease associations and has been a most common used storage of miRNA functional similarity (Li *et al.*, 2022; Zhou *et al.*, 2021). Here we extracted the complete data set from the website and then converted the pri-miRNA to mature miRNA using ID-cross file derived from miRBase (https://mirbase.org/). A miRNA network containing 375136 edges and 867 nodes was obtained.

**2.5 mRNA similarity network**

A comprehensive network backing on six distinct evidence types of orthology-based model organism, gene fusion events, gene proximity, co-expression, experiments and database was downloaded from STRING v12.5 (https://string-db.org). According to the original paper of STRING database (Szklarczyk *et al.*, 2021), experimentally supported evidences were adopted to ensure the reliability of data. The final mRNA similarity network was obtained by the Bayesian network integration approach provided by STRING using python. 5875978 edges between19113 mRNAs were constructed.

**2.6 Highlighting regulatory elements in heterogeneous network**

Heterogeneous networks were ensembled by linking independent similarity networks with their crosstalks, which were mainly derived from experimentally verified regulatory data. The crosstalks between circRNA and miRNA similarity network, lncRNA and miRNA similarity network, miRNA and mRNA similarity network, lncRNA and mRNA similarity network were the circRNA-miRNA, lncRNA-miRNA, miRNA-mRNA, lncRNA-mRNA regulations respectively. If one mRNA and one circRNA shared the same miRNA targets, they were linked between circRNA and mRNA similarity network. If miRNA's targets were enriched in the pathway by hypergeometric test (Pian *et al.*, 2020), then miRNA-pathway crosstalks were determined, which linked pathway similarity network and miRNA similarity network. Pathway similarity network and mRNA similarity network were connected by mRNA members of a pathway. The diffusion algorithm, RWR-MH (Valdeolivas *et al.*, 2019; Ran *et al.*, 2021) was adapted to highlight regulatory elements by reducing the noises of heterogeneous and measuring the similarity of biological elements in the same low-dimensional space.

#### S3 Application

The cellular senescence, driven by interconnected molecular changes, has turned into a significant risk role in highly prevalent chronic and devastating diseases, including neurodegeneration, cardiovascular disorders, and cancer, as widely demonstrated in previous research (López-Otín *et al.*, 2013; Marengoni *et al.*, 2011). Here, through a compiled cellular senescence-related miRNA and mRNA mixed data set, we performed a regular workflow for tentatively screening candidate targets with NetLCP. Totally 321 mRNAs from the Aging Atlas database (https://ngdc.cncb.ac.cn/aging) and CellAge (https://genomics.senescence.info/cells) and 48 miRNAs from a review article (Munk *et al.*, 2017) were derived respectively.

In this example, we first highlighted regulatory elements (lncRNA, circRNA and pathway) (3.1) with NetLCP. The top 10 of highlighted lncRNAs and pathways (KEGG, Reactome and Wikipathway) and the input transcriptome were combined to explore their combinations of regulatory elements (CREs) in the local area of heterogenous network. Here we concentrated on lncRNA-miRNA-mRNA-pathway regulation. Furthermore, we prioritized the CREs by the number of their eQTLs (3.2). Ultimately, we revealed the variant 'switches' i.e. variants on binding sites in selected prioritized CREs. (3.3).

##### 3.1 Highlighting the regulatory elements

We first highlighted the regulatory elements and illustrated the top 10 prioritizations (the disease pathways were excluded) with the senescence-associated evidences by text validation in PubMed (see **Table 1**). The evidences were categorized into two types, of which R representing the element had been demonstrated to be related to cellular senescence, and B indicating that there was no direct link to cellular senescence but the biological associations were revealed in text validation. In the top result of each heterogeneous network, hsa04218 (cellular senescence) and R-HSA-2559580 (Oxidative Stress Induced Senescence) were the well-defined senescence pathway. WP382 (MAPK signaling pathway) were demonstrated to be a hall marker of cellular senescence. The circRNA hsa_circ_0001006 and lncRNA XIST could involve in cellular senescence by regulating hsa-miR-214-3p (Zhong *et al.*, 2022).

**Table1. Top 10 highlighted elements in the five heterogeneous networks**

| KEGG | pathID | FunScore | Empirical p.value | Evidence | PMID |
|---|---|---|---|---|---|
| **Cellular senescence** | hsa04218 | 2.180 | 7.69E-113 | B / R | ● |
| **PI3K-Akt signaling pathway** | hsa04151 | 1.747 | 1.11E-91 | B / R | 35505269 |
| **Thyroid hormone signaling pathway** | hsa04919 | 1.727 | 4.68E-71 | B / R | 32360904 |
| **FoxO signaling pathway** | hsa04068 | 1.722 | 2.23E-92 | B / R | 35154571 |
| **HIF-1 signaling pathway** | hsa04066 | 1.712 | 4.25E-61 | B / R | 34385092 |
| **MAPK signaling pathway** | hsa04010 | 1.629 | 3.68E-89 | B / R | 35358003 |
| **Signaling pathways regulating pluripotency of stem cells** | hsa04550 | 1.619 | 5.88E-64 | B / R | 34519176 |
| **Cell cycle** | hsa04110 | 1.363 | 3.63E-90 | B / R | 34667279 |
| **Sphingolipid signaling pathway** | hsa04071 | 1.333 | 1.58E-50 | B / R | 34069611 |
| **Rap1 signaling pathway** | hsa04015 | 1.309 | 3.22E-66 | B / R | 31742863 |
| **Reactome** | **pathID** | **FunScore** | **Empirical p.value** | **Evidence** | **PMID** |
| **Oxidative Stress Induced Senescence** | R-HSA-2559580 | 5.628 | 4.78E-51 | B / R | ● |
| **Transcriptional Regulation by E2F6** | R-HSA-8953750 | 4.593 | 3.50E-34 | B | 10727462 |

| | | | | | |
|---|---|---|---|---|---|
| **Cyclin E associated events during G1/S transition** | R-HSA-69202 | 4.027 | 8.52E-16 | B | 31112603 |
| **Oncogene Induced Senescence** | R-HSA-2559585 | 3.920 | 7.28E-29 | B / R | ● |
| **Regulation of PTEN gene transcription** | R-HSA-8943724 | 3.814 | 1.30E-17 | B / R | 34643308 |
| **SUMOylation of DNA damage response and repair proteins** | R-HSA-3108214 | 3.622 | 5.26E-16 | B / R | 35349763 |
| **SUMOylation of chromatin organization proteins** | R-HSA-4551638 | 3.559 | 2.14E-26 | B / R | 35338388 |
| **ISG15 antiviral mechanism** | R-HSA-1169408 | 3.406 | 5.45E-18 | B | 33555479 |
| **SUMOylation of RNA binding proteins** | R-HSA-4570464 | 3.048 | 8.42E-16 | B | 35216455 |
| **SUMOylation of DNA methylation proteins** | R-HSA-4655427 | 3.032 | 3.29E-14 | B/R | 35371604 |
| **CircRNA** | **Evidence** | **FunScore** | **Empirical p.value** | **Total In PubMed** | **PMID** |
| **hsa_circ_0001006** | B | 3.417 | 2.23E-214 | 1 | 35306484 |
| **hsa_circ_0000677** | B | 3.206 | 1.79E-192 | 1 | 31960989 |
| **hsa_circ_0001417** | B | 3.078 | 3.88E-156 | 1 | 32808350 |
| **hsa_circ_0000005** | B | 3.078 | 1.45E-179 | 1 | 33663506 |
| **hsa_circ_0001105** | NA | 2.972 | 1.43E-151 | 0 | |
| **hsa_circ_0000284** | B | 2.887 | 4.10E-155 | 5 | 34135597 |
| **hsa_circ_0000006** | B | 2.867 | 4.94E-121 | 1 | 34532379 |
| **hsa_circ_0033351** | NA | 2.803 | 1.49E-149 | 0 | |
| **hsa_circ_0001756** | B | 2.798 | 1.45E-123 | 1 | 35113003 |
| **hsa_circ_0023117** | NA | 2.773 | 5.46E-95 | 0 | |
| **Wikipathway** | **pathID** | **FunScore** | **Empirical p.value** | **Evidence** | **PMID** |
| **MAPK signaling pathway** | WP382 | 3.344 | 4.25E-37 | B / R | 35358003 |
| **VEGFA-VEGFR2 signaling pathway** | WP3888 | 3.205 | 3.89E-46 | B | 32300208 |
| **Senescence and autophagy in cancer** | WP615 | 2.898 | 5.64E-42 | B / R | ● |

| | | | | | |
|---|---|---|---|---|---|
| **TGF-beta signaling pathway** | WP366 | 2.554 | 1.92E-40 | B / R | 35305149 |
| **Adipogenesis** | WP236 | 2.380 | 1.77E-20 | B | 34712331 |
| **Insulin signaling** | WP481 | 2.278 | 3.12E-15 | B | 35154571 |
| **PI3K-Akt signaling pathway** | WP4172 | 2.274 | 2.13E-34 | B / R | 35361048 |
| **Cell cycle** | WP179 | 2.243 | 1.40E-16 | B / P | |
| **DNA damage response** | WP707 | 2.235 | 1.35E-31 | B / P | 35365245 |
| **EGFR tyrosine kinase inhibitor resistance** | WP4806 | 2.16 | 5.94E-15 | B | 32323422 |

| **LncRNA** | **Ensembl ID** | **FunScore** | **Empirical p.value** | **Evidence** | **PMID** |
|---|---|---|---|---|---|
| **XIST** | ENSG00000229807 | 5.809 | 6.23E-4 | B | 35196590 |
| **OIP5-AS1** | ENSG00000247556 | 5.378 | 1.24E-4 | B / R | 31476304 |
| **MALAT1** | ENSG00000251562 | 5.117 | 1.551E-4 | B / R | 35309907 |
| **FGD5-AS1** | ENSG00000225733 | 4.811 | 1.477E-4 | B | 32675387 |
| | ENSG00000230551 | 4.663 | 3.054E-3 | B | 26975529 |
| **HCG18** | ENSG00000231074 | 4.474 | 6.91E-4 | B | 35389764 |
| **KCNQ1OT1** | ENSG00000269821 | 4.223 | 5.454E-3 | B | 35218109 |
| **HCG17** | ENSG00000270604 | 4.222 | 2.397E-3 | NA | |
| **SNHG1** | ENSG00000255717 | 4.055 | 5.80E-3 | B | 34618681 |
| **GAS5** | ENSG00000234741 | 3.941 | 2.659E-2 | B / R | 34386495 |

*• represents the well-defined relationship with senescence.

##### 3.2 Prioritizing CREs

Furthermore, we combined the top 10 elements of highlighted lncRNAs and pathways (KEGG, Reactome and Wikipathway) with the input transcriptome, and then used NetLCP to explore their CREs in the local area of heterogenous network. **Supplementary Fig.1** exhibited the overview of CRE network and the degree distribution of nodes. Obviously, miRNA elements (colored by red) could be regarded as the main mediums in the whole regulation since they held the largest proportions of degree in the network. Overall, among three types of elements, MIMAT0000255 (hsa-miR-34a-5p), 4193 (MDM2), ENSG00000231074 (HCG18), R-HSA-2559582 (Senescence-Associated Secretory Phenotype) and hsa04151/WP4172 (PI3K-Akt signaling pathway) took the largest node degrees, i.e. 107, 48, 49, 19, 41/41 respectively.

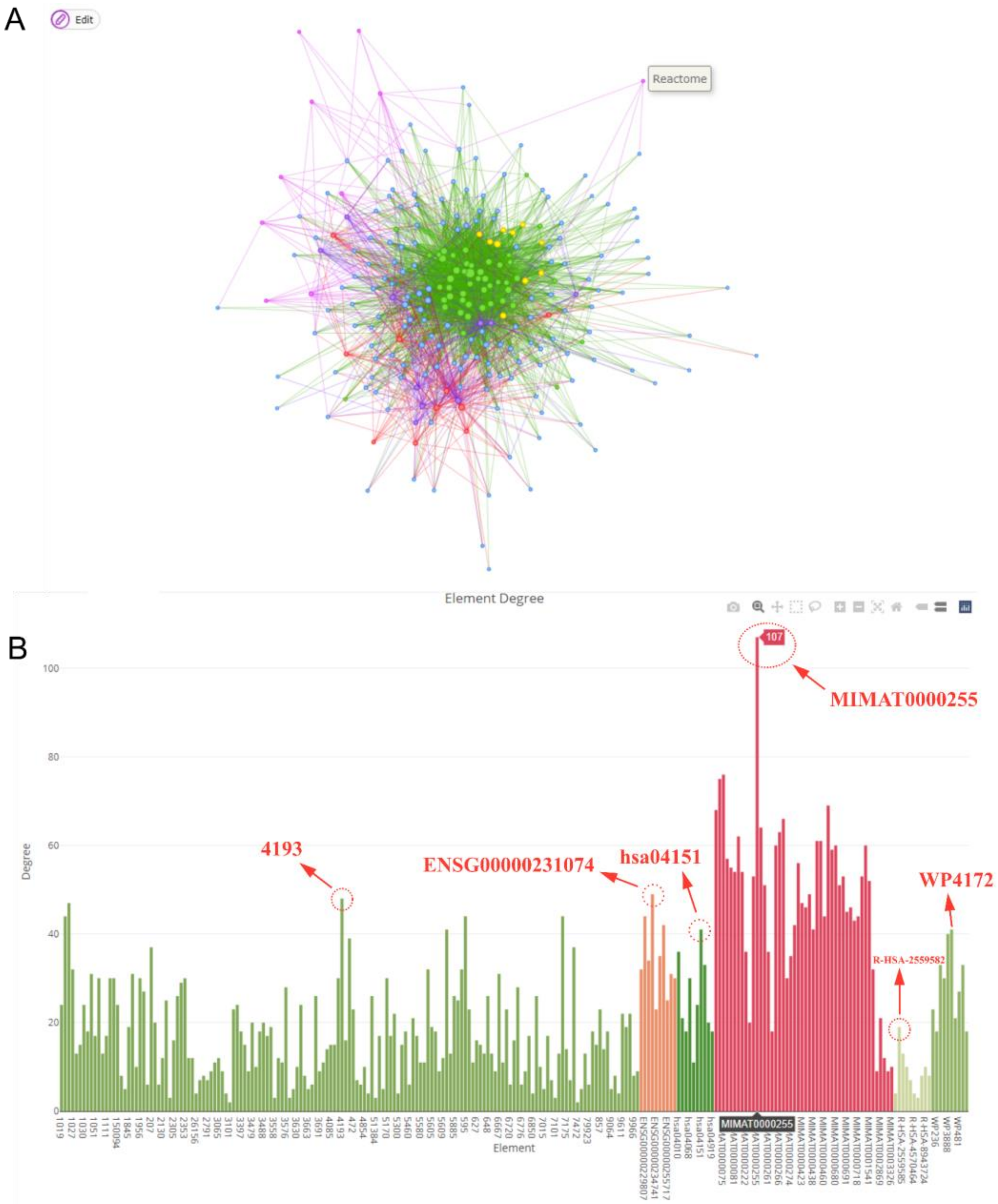


**Supplementary Fig.1 Overview of CREs in local area of heterogeneous network A) Network visualization of CREs B) Node distribution of network**

Then we extracted a subnetwork by selecting the elements with their degrees ≥ 40. The subnetwork was showed in **Supplementary Fig.2A**. Obviously, the CREs containing 4193 (MDM2) were regulated by the most eQTLs (**Supplementary Fig.2B**). Therefore, we selected them to explore the variant ‘switches’ of CREs in the local area of heterogeneous network.

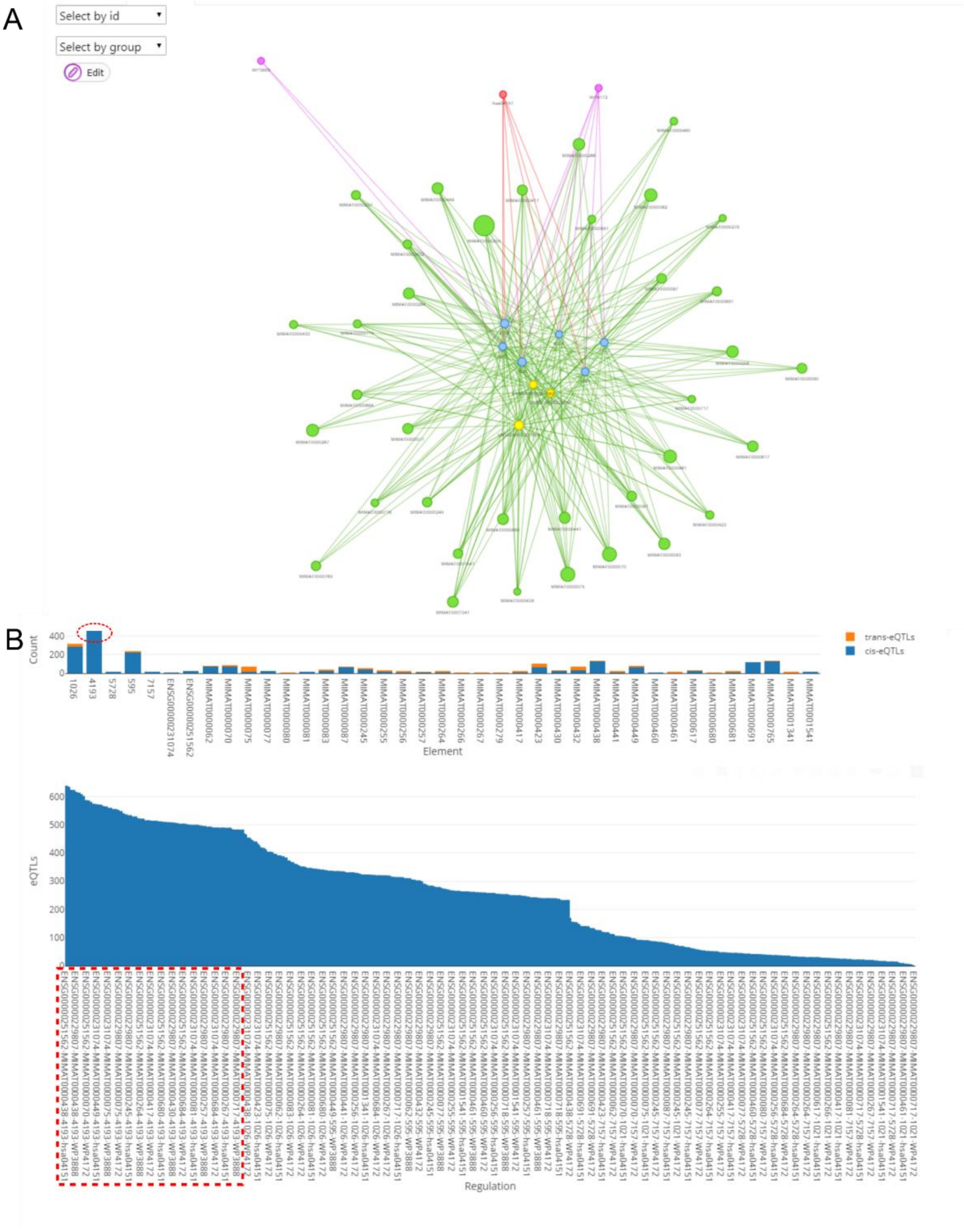

**Supplementary Fig.2 A) Subnetwork visualization of CREs B) The distribution of eQTLs of CREs in subnetwork**

##### 3.3 Detecting variant 'switches' in the top prioritized CREs

The variant 'switches'detected on the binding site of selected CREs containing 4193(MDM2) (3.2) were summarized in **Supplementary Fig.3**. The CREs ENSG00000251562(MALAT1)→MIMAT0000081(hsa-miR-25-3p)→4193(MDM2)→hsa04151(PI3K-Akt signaling pathway)/WP3888(VEGFA-VEGFR2 signaling pathway)/WP4172(PI3K-Akt signaling pathway) were highlighted. The expression of MALAT1 was significantly increased in senescent neuroblastoma cells (Zhang *et al.*, 2022). The hsa-miR-25-3p adversely affected apoptotic cell death, cell cycle arrest and cellular senescence (Kumar *et al.*, 2011). MDM2 promoted cellular senescence by modulating WRN stability (Liu *et al.*, 2019). NetLCP integrated their variant 'switches' into CRE network to intuitively exhibit the regulatory information and also performed statistics.

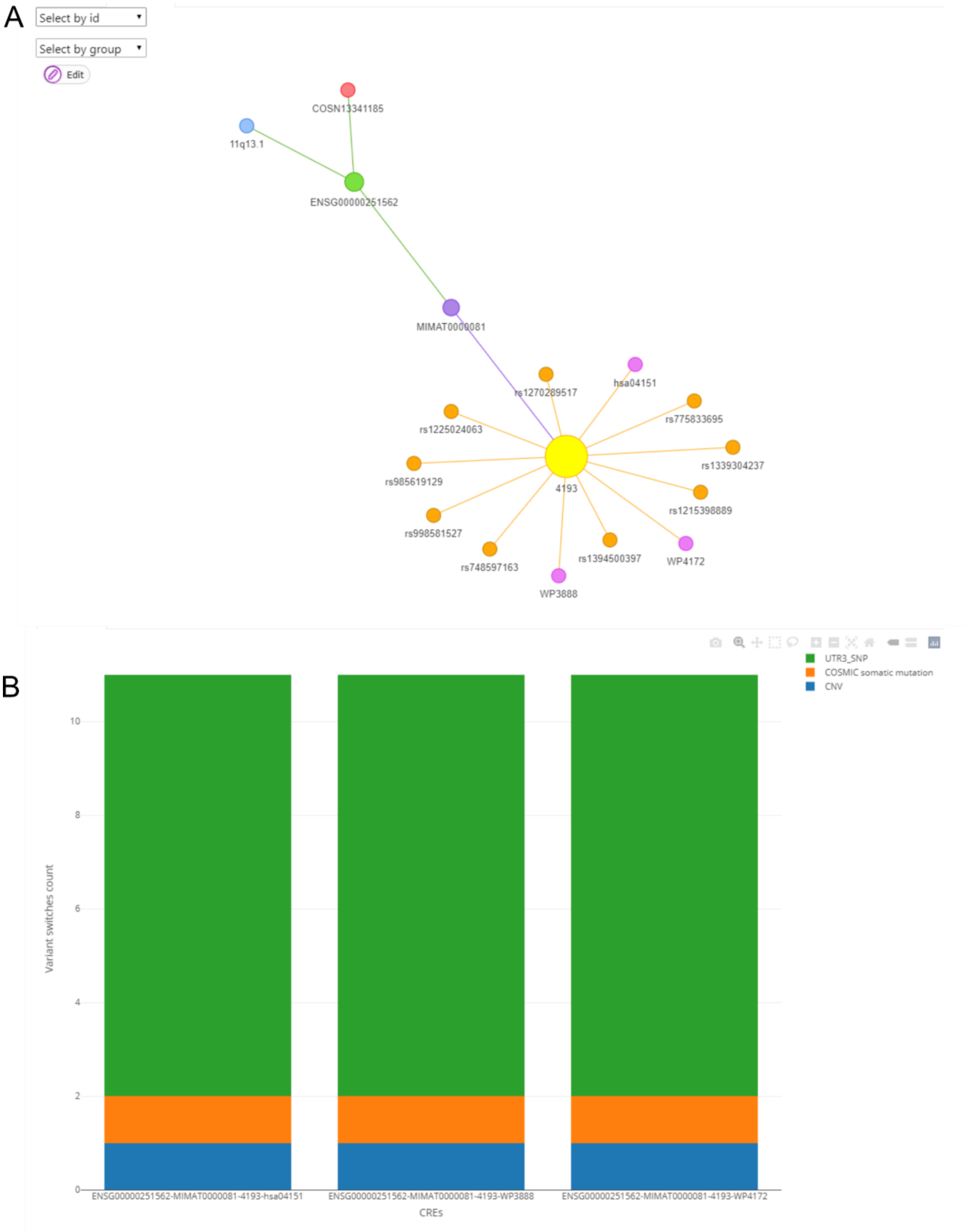


**Supplementary Fig.3 The regulatory variants on the binding site of top CREs A) variant 'switches' network B) variant 'switches' statistics**

##### 3.4 Application with NetLCP GUI

Launch NetLCP GUI by the command NetLCPGUI () in R language.

###### Highlighting regulatory elements

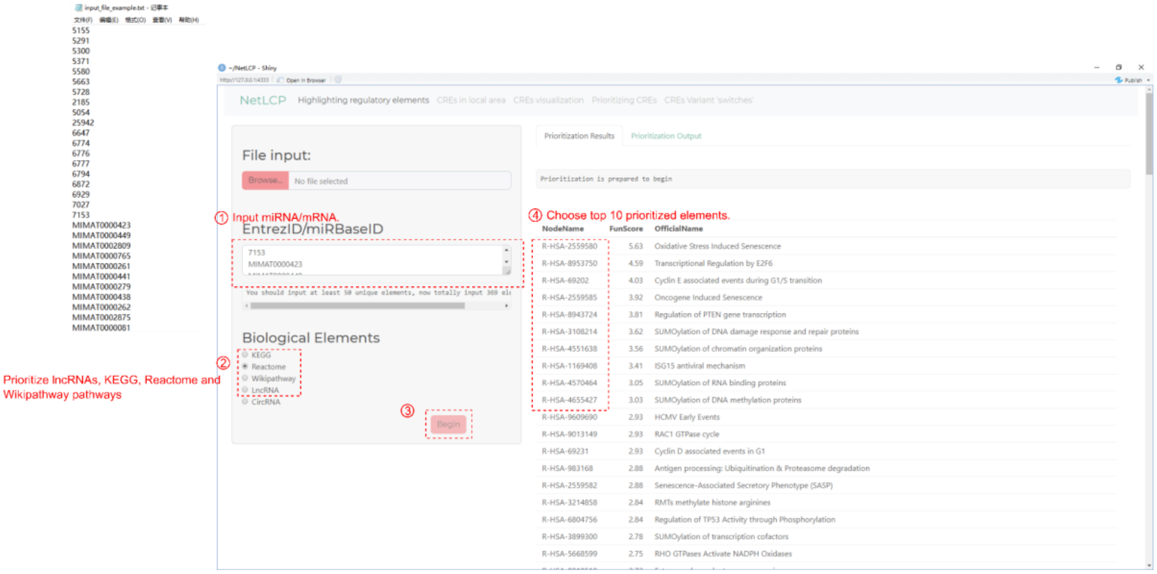


###### Combining top 10 highlighted regulatory elements with input miRNA/mRNA to explore the CREs

Explore CREs

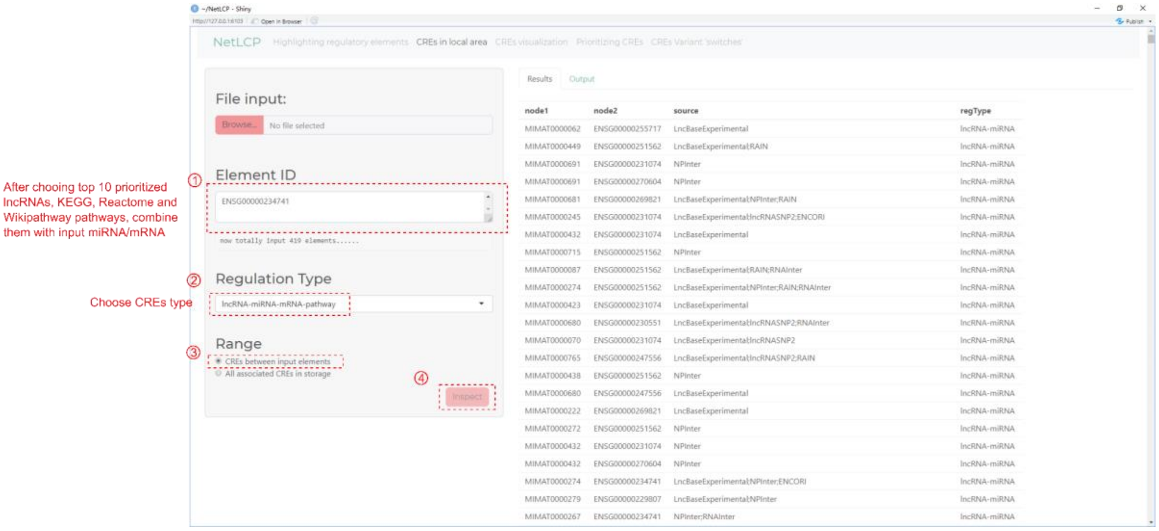

Statistics of node degree in CREs network

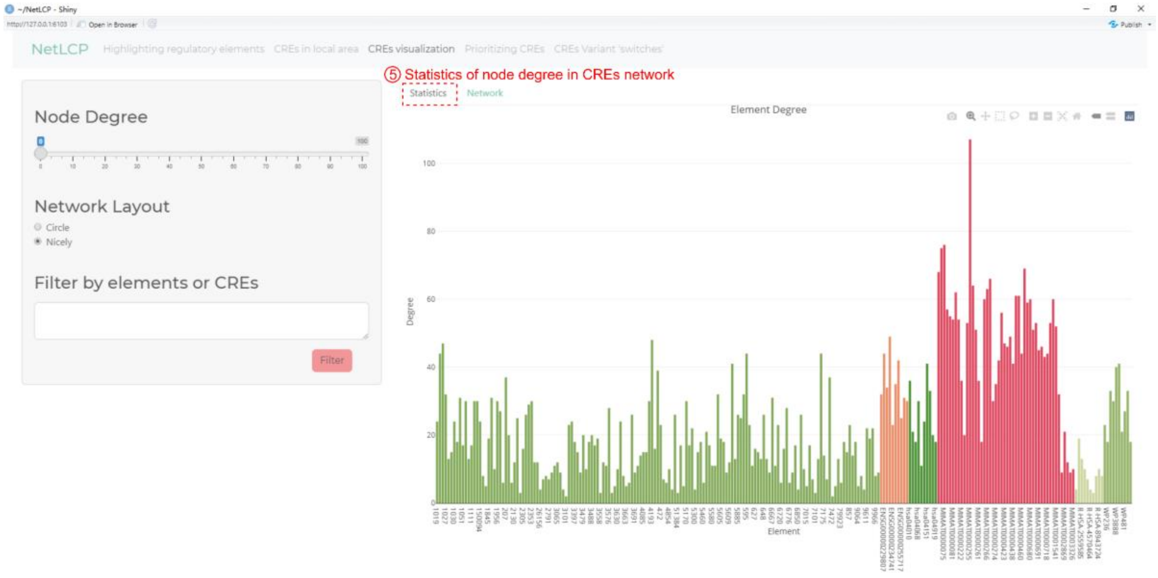


Network visualization of CREs

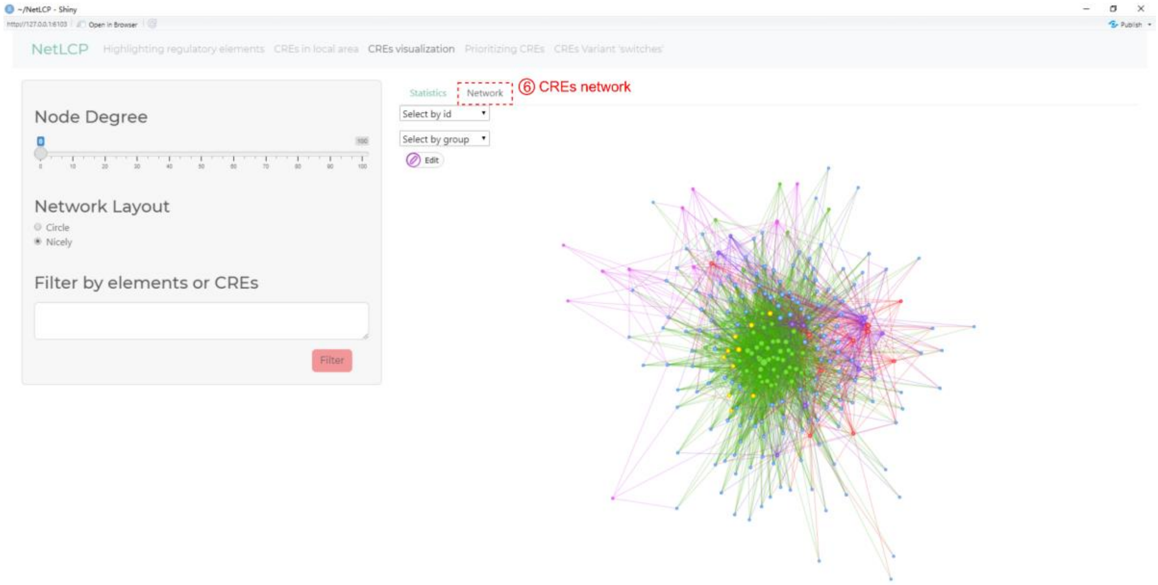

### Prioritizing CREs

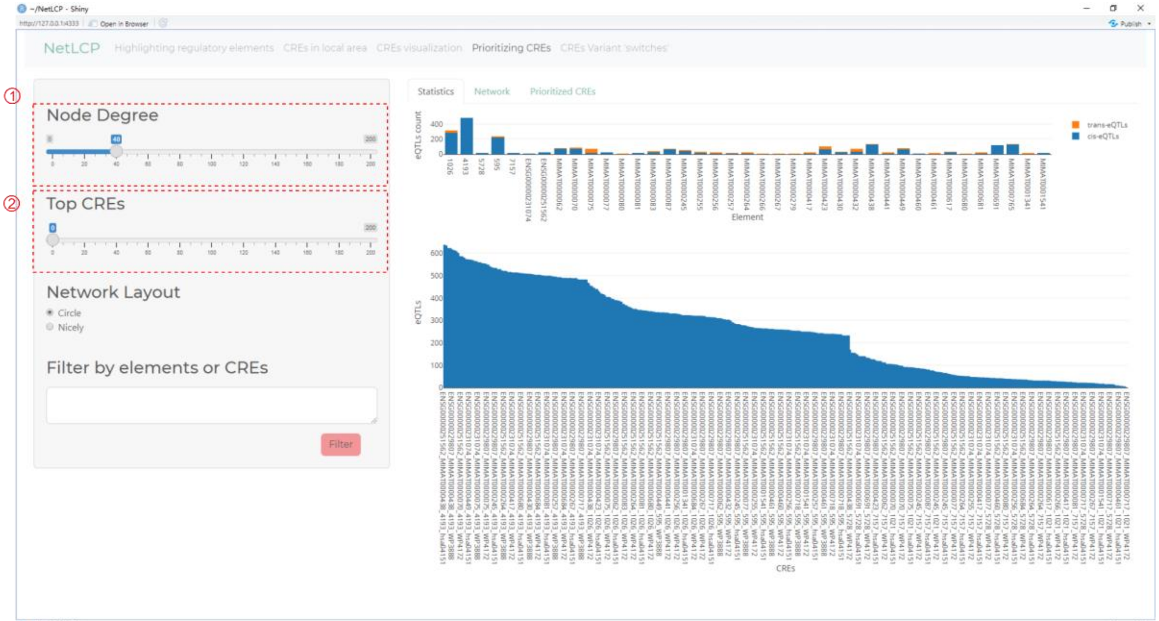


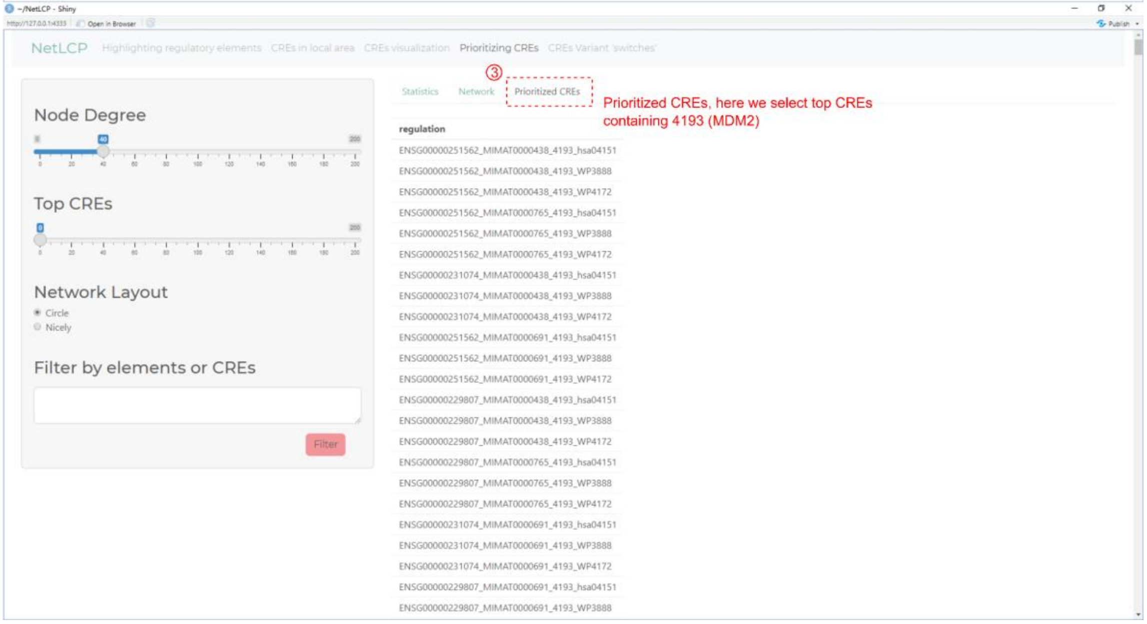

#### Exploring the variant 'switches' in top prioritized CREs

Statistics

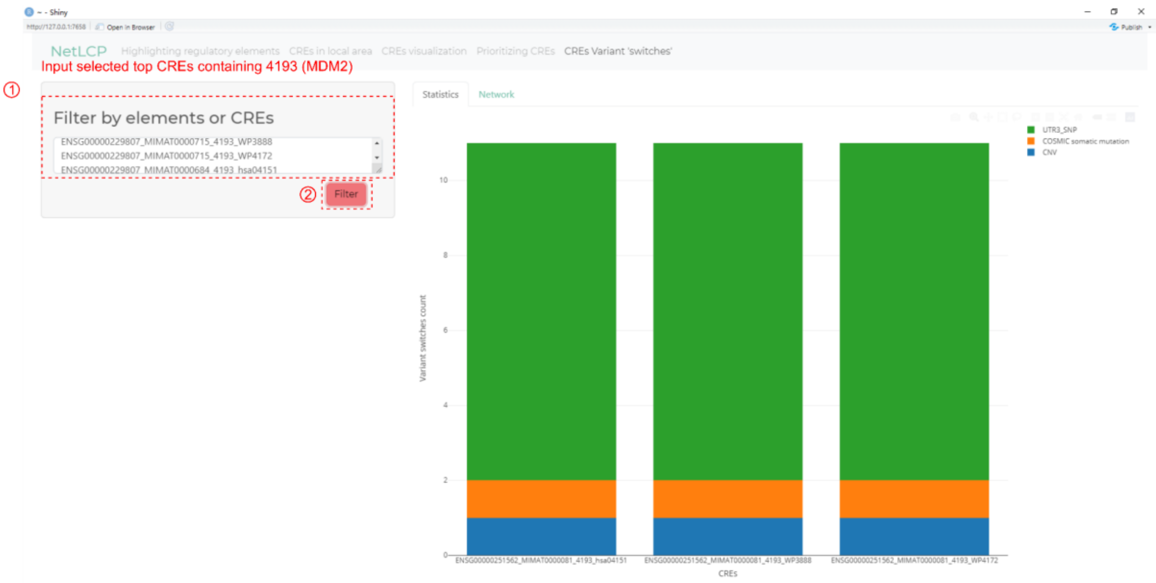

Network visualization

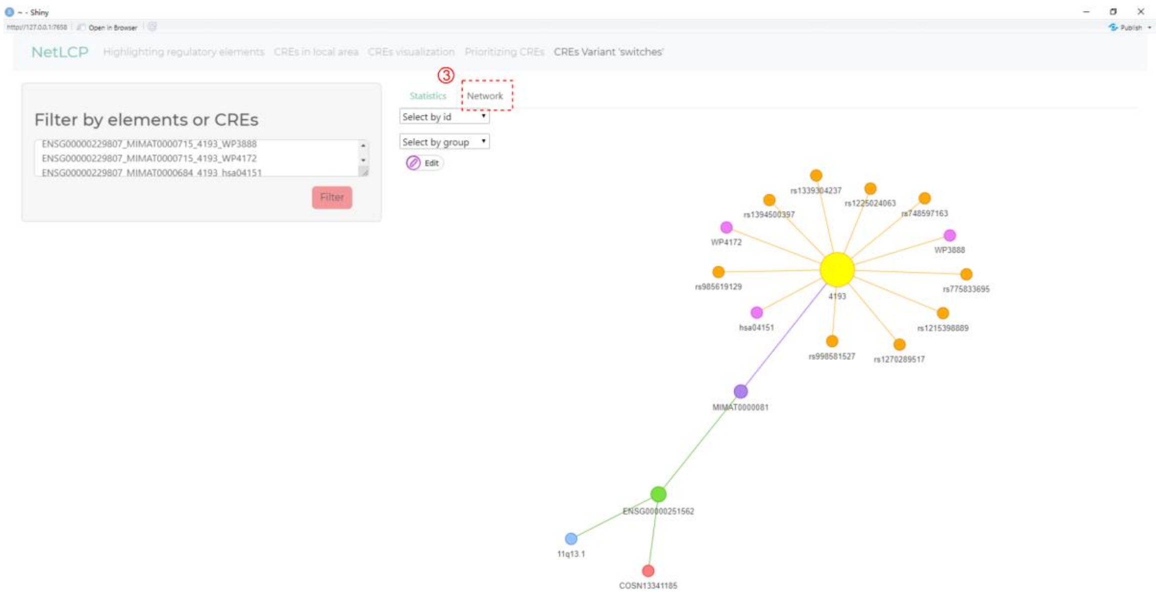

#### S4 Tutorial

NetLCP command and GUI tutorial can be found at https://github.com/mortyran/NetLCP.
